# Defining Cardiac Cell Populations and Relative Cellular Composition of the Early Fetal Human Heart

**DOI:** 10.1101/2021.10.21.465281

**Authors:** Jennifer M. Dewing, Vinay Saunders, Ita O’Kelly, David I. Wilson

## Abstract

The human heart is primarily composed of cardiomyocytes, fibroblasts, endothelial and smooth muscle cells. Reliable identification of fetal cardiac cell types using protein markers is important for understanding cardiac development and delineating the cellular composition of the human heart during early development, which remains largely unknown. The aim of this study was to use immunohistochemistry (IHC), flow cytometry and RT-PCR analyses to investigate the expression and specificity of commonly used cardiac cell markers in the early human fetal heart (8-12 post-conception weeks). The expression of previously reported protein markers for the detection of cardiomyocytes (Myosin Heavy Chain (MHC) and Troponin I (cTnI)), fibroblasts (DDR2, Thy1, Vimentin), endothelial cells (CD31) and smooth muscle cells (α-SMA) were assessed. Flow cytometry revealed two distinct populations of cTnI expressing cells based on fluorescence intensity: cTnI^High^ and cTnI^Low^. MHC positive cardiomyocytes were cTnI^High^, whereas MHC negative non-myocyte cells were cTnI^Low^. cTnI expression in non-myocytes was further confirmed by IHC and RT-PCR analyses, suggesting troponins are not cardiomyocyte-specific and may play distinct roles in non-muscle cells during early development. Vimentin was confirmed to be enriched in cultured fibroblast populations and flow cytometry revealed Vim^High^ and Vim^Low^ cell populations in the fetal heart. MHC positive cardiomyocytes were Vim^Low^ whilst CD31 positive endothelial cells were Vim^High^. Based on the markers investigated, we estimate fetal human cardiomyocyte populations comprise 75-80% of total cardiac cells and exhibit the following marker profile: α-MHC^+^/cTnI^High^/Vim^Low^. For the non-cardiomyocyte population, we estimate they comprise 20-25% of total cardiac cells and exhibit the following marker profile: α-MHC^-^/cTnI^Low^/Vim^High^. Our study suggests the marker profiles and proportions of fetal cardiac populations are distinct from that of the adult heart.

## Introduction

The human heart is primarily composed of cardiomyocytes, fibroblasts, endothelial and smooth muscle cells. Rodent studies have proposed that cardiomyocytes occupy ∼75% of normal adult myocardial volume and account for 30-40% of cell number, with non-cardiomyocytes (endothelial, smooth muscle cells and fibroblasts), though smaller in size, being the predominant cell type^1-5^. Similarly, immunohistochemical and flow cytometry analysis by Pinto *et al*^4^ concluded that the adult human heart consists of approximately 30% cardiomyocytes and 50% endothelial cells. The relative cellular composition of the adult heart is likely to differ to that of the fetal heart during the dynamic stages of early development. The identification, characterisation and purification of cardiac cells relies upon their expression of cell-specific protein markers and is key to better understanding cardiac development and delineating the cellular composition of the human heart during early development, which remains largely unknown. Furthermore, given that pluripotent stem cell-derived cardiomyocytes (PSC-CMs) have been shown to be phenotypically similar to cardiomyocytes of the mid-gestation human fetal heart, an improved understanding of the protein marker profiles of these cells could aid the differentiation and purification of PSC-CMs in regenerative medicine^6^.

One of the key challenges to the isolation and identification of cell populations in both adult and fetal hearts is the heterogeneity of fibroblast cells and the lack of a defined cardiac fibroblast-specific marker. Fibroblasts are cells of mesenchymal origin that produce extracellular matrix proteins including collagen and fibronectin ^7^. The cell surface collagen receptor DDR2 has previously been used as a marker of cardiac fibroblasts ^8-11^, however, its expression has also been reported in rat endothelial and smooth muscle cells, although absent from cardiomyocytes ^2,10,12,13^, raising the possibility of interspecies differences. The cytoskeletal intermediate filament vimentin is a commonly used fibroblast marker due to its high levels of expression in cells of mesenchymal origin. However, vimentin expression in endothelial and smooth muscle cells underscores this protein’s lack of specificity^14,15^. Thy-1 (CD90), a cell-surface glycoprotein, has been shown to be expressed on cultured cardiac fibroblasts and as a surface protein it has the advantage over vimentin and DDR2 in that it does not require cell permeabilisation and can therefore be used for fluorescent activated cell sorting (FACS) ^16-18^. Thy-1 has been detected in cultured rat cardiac fibroblasts, with increased expression detected in fibrotic areas within the rat heart, suggesting it may be a marker for proliferating fibroblasts ^16^. However, Thy-1 has also been detected in thymocytes, T-cells, neurons, hematopoietic stem cells and endothelial cells ^19-21^. Whilst Pinto *et* al., were able to use Thy-1 and Sca-1 to delineate cardiac fibroblasts in mice, neither of these markers were successful at isolating cardiac fibroblasts of the adult human heart^4^. Similarly, fibroblast-specific protein 1 (FSP1) has been shown to lack fibroblast specificity in cardiac tissue during remodelling, with expression identified in hematopoietic, endothelial and vascular smooth muscle cells ^22^. Together, these studies suggest that many commonly used markers of fibroblasts lack specificity.

Endothelial and smooth muscle cells that line the cardiac vasculature can be detected by their expression of CD31 (PECAM-1) and α-smooth muscle actin (α-SMA), respectively. Commonly used markers of cardiomyocytes include the sarcomeric proteins cardiac troponin (subunits I, T and C), myosin and tropomyosin. α-myosin heavy chain (α-MHC) has high expression levels in cardiac muscle and significantly lower levels in skeletal muscle; subsequently the α-MHC promoter is often used in transgenic mouse models to track cardiomyocytes ^18,23,24, 25^. Whilst these markers may identify mature, differentiated endothelial, smooth muscle and cardiomyocyte cell populations in adult tissue, their specificity during development may be distinct.

The aim of this study was to use immunohistochemistry (IHC) and flow cytometry, alongside RT-PCR analysis to evaluate the specificity of a range of markers that hold the potential to specifically define cell populations in the early human fetal heart and use the novel marker profiles to estimare the cellular composition of the heart during early development.

## Materials and Methods

### Isolation of Cardiac Cells

Human fetal heart tissue (8 to 12 post conception weeks (pcw) was obtained from the Human Developmental Biology Resource (HDBR) Newcastle, UK. Tissue collection was in agreement with the Declaration of Helsinki (ethics approval reference: 08/HO906/21+5 NRES Committee North East-Newcastle & North Tyneside). Heart tissue was dissected to remove the aorta and pulmonary artery and the remaining ventricles and atria were divided into 1mm^3^ pieces using a tissue chopper (McIlwain) and mechanically dissociated together in 3ml of phosphate buffered saline (PBS) using the gentleMACS dissociator machine (Miltenyl Biotec) using the programme pre-set for heart tissue. The cell suspension was passed through a 70 μm nylon cell strainer (BD Biosciences). For analysis of cells from the aorta and pulmonary artery, the process was repeated with dissected aorta and pulmonary artery tissue.

### Fibroblast Isolation

Fibroblasts were isolated from human fetal heart and skin tissue by explant migration following previously reported methods by Ieda *et al*., (2010)^18^. Briefly, tissue was chopped into 1mm^3^ pieces using a tissue chopper (McIlwain) and cultured in 10% FBS DMEM on gelatin-coated plates for 1 to 2 weeks to facilitate fibroblast migration. The fibroblasts cells were then isolated by addition of trypsin followed by filtering through a 70 μm filter to allow single fibroblast cells to pass through. The cellular filtrate was then pelleted by centrifugation at 1300 rpm for 3 minutes and re-plated.

### Flow Cytometry

Dissociated and filtered cardiac cells were further filtered through a 35 μm cap into FACS tubes (BD Biosciences) to ensure a single cell suspension. Cells were treated with human FcR blocking serum (BD Bisociences), fixed in 4% formaldehyde (Sigma) and permeabilized in 0.05% saponin (Sigma). Cells were incubated with primary and secondary antibodies and analysed using a BD FACSCanto I Flow cytometer (BD biosciences). Single, dual and triple stain flow cytometry analyses were carried out. Positive cells were gated based on their expression above isotype and no primary control samples.

### Immunohistochemistry and Immunocytochemistry

Immunohistochemistry followed previously reported methods^26^. Briefly, human fetal heart tissue was fixed in 4% formaldehyde, embedded in paraffin and 5 μm microtome sections cut. Antigen retrieval was performed (15 minutes of boiling in 0.1mM sodium citrate) before primary and secondary antibody incubations and 4’, 6-diamidino-2-pheylindole (DAPI) staining. For immunocytochemistry of cultured primary cardiac fibroblasts, cells were cultured on glass coverslips in 6-well plates and fixed in 4% formaldehyde. Cells were blocked with 3% BSA in PBS and treated with primary antibodies diluted in 0.1% triton overnight at 4 °C, followed by 1hr with secondary antibodies diluted in 0.1% triton.

### Antibodies for Flow Cytometry and Immunohistochemistry

Antibodies were selected to identify cardiomyocytes, fibroblasts, smooth muscle cells and endothelial cells, based on current literature ^9,10,16,25,27^. These included: Myosin Heavy chain (MHC) conjugated to efluor660 (50-6503-80 ebioscience), Troponin I (Ab47003, Abcam), DDR2 (Sc7555 Santa Cruz), Thy-1, (555595 BD Biosciences), Vimentin conjugated to Cy3 (C9080 Sigma), CD31 to APC (21270316 Immunotools) and α-SMA conjugated to FITC (53-9760-80 ebioscience). Appropriate isotype control antibodies were used as negative controls for directly conjugated antibodies, and a no primary antibody control was used for non-conjugated antibodies.

### RT-PCR

Total RNA was isolated using TRIzol Reagent (Invitrogen) from whole heart tissue and cell pellets of cultured primary fibroblasts. 1 μg of RNA was used for cDNA synthesis using M-MLV reverse transcriptase (Promega). Primer sequences and amplicon sizes are listed in supplementary table 1. Polymerase Chain Reaction (PCR) consisted of initial denaturation of DNA at 95 °C, followed by 40 cycles of 94 °C for 1 minute, annealing at 58 °C for 1 minute and extension at 72 °C for 1 minute, with a final extension of 72 °C for 10 minutes.

**Supplementary Table 1.**
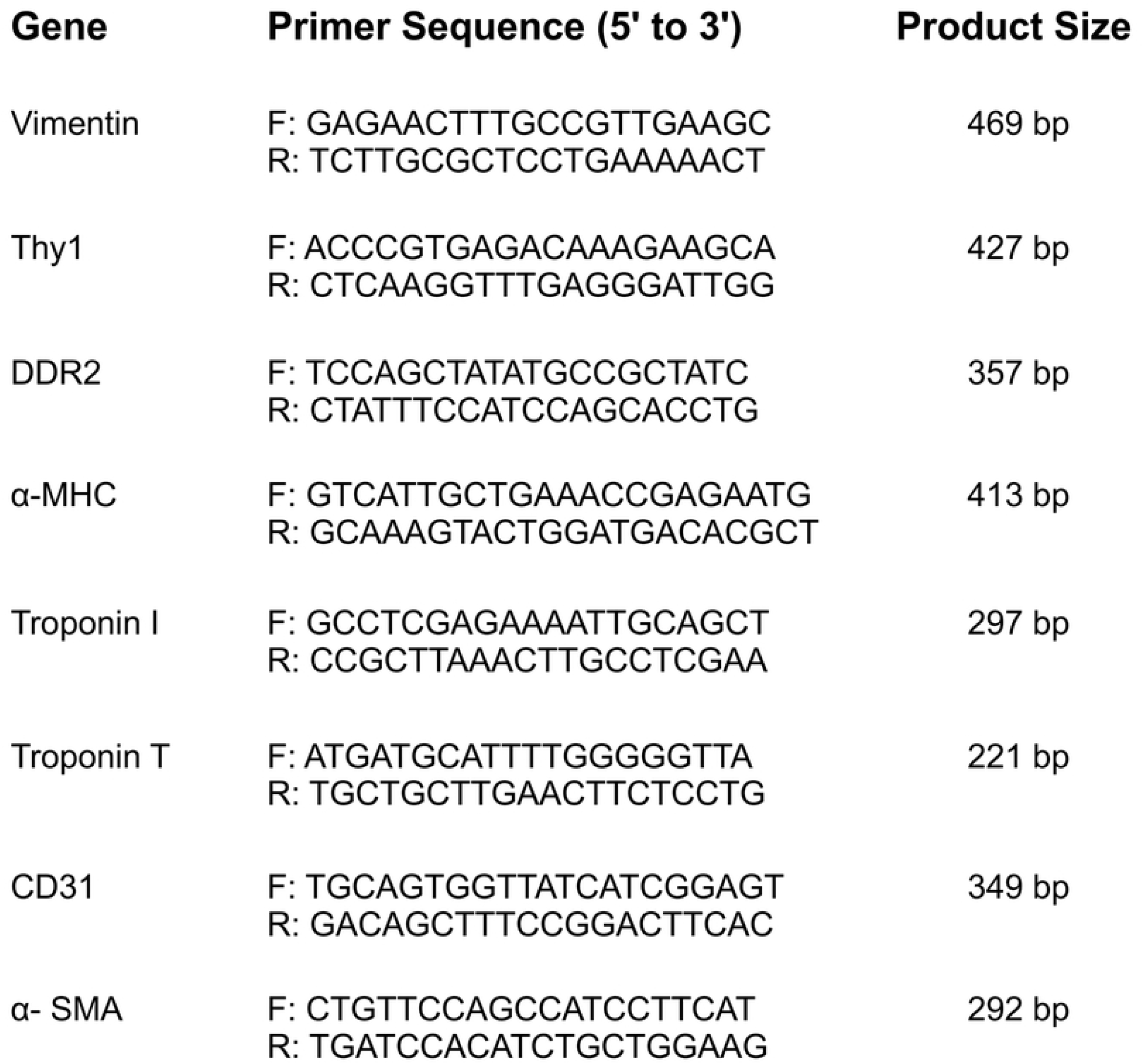
Primer sequences and amplicon sizes Primer sequences used for the detection of mRNA transcripts of key cell markers using RT-PCR.

## Results

### Single marker immunohistochemistry and flow cytometry of the fetal human heart

Fetal human heart tissue was stained for markers of cardiomyocytes (MHC, TnI, TnT), cardiac fibroblasts (vimentin, DDR2, Thy-1), endothelial cells (CD31) and smooth muscle cells (α-SMA) (Figure 1). TnI and α-MHC showed distinct sarcomeric staining typical of cardiomyocytes (Fig 1A). The fibroblast markers showed extensive staining throughout the heart tissue, with vimentin and Thy-1 staining localised predominantly to the cytoplasm (Fig 1B). CD31 and α-SMA showed positive staining of the endothelial and smooth muscle cells lining the vessels in the heart, respectively (Fig 1C).

**Figure 1.**
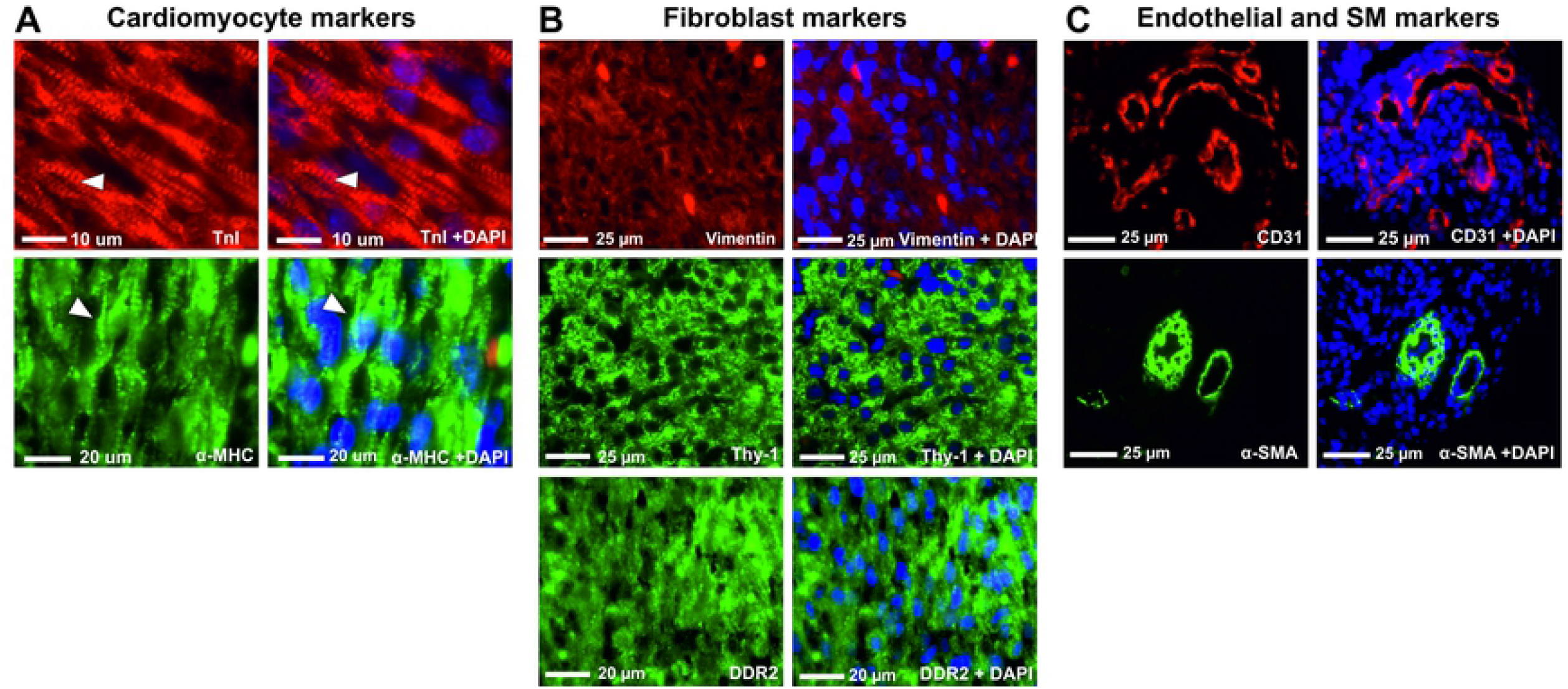
Immunohistochemistry of fetal human heart tissue stained for cardiac cell markers. (A) Immunohistochemistry of cardiomyocyte markers TnI and α-MHC. (B) Immunohistochemistry of fibroblast markers vimentin, Thy-1 and DDR2. (C) Immunohistochemistry of endothelial and smooth muscle markers, CD31 and α-SMA, respectively. DAPI was used as a counter stair for cell nuclei.

Expression of these markers in dissociated cardiac cells, detected by flow cytometry, was used to inform on the proportions of the major cells types of the fetal human heart (Fig 2). For cardiomyocyte marker expression, 75% of cells were MHC^+^ (n=21, SEM ±1.40) (Fig 2B) and 93% were TnI^+^ (n=14, SEM ±1.5) (Fig 2C). For cardiac fibroblast markers, 90% of cells were vimentin^+^ (n=18, SEM ±1.2) (Fig 2D), 83% were DDR2^+^ (n=4, SEM ±0.6) (Fig 2E) and 81% were Thy-1^+^ (n=15 SEM ±1.40) (Fig 2F). For endothelial and smooth muscle cell markers, 9% of cells were CD31^+^ (n=12, SEM= ±0.54) (Fig 2G) and 12% were α-SMA^+^ (n=7, SEM= ± 1.9) (Fig 2H), respectively.

**Figure 2.**
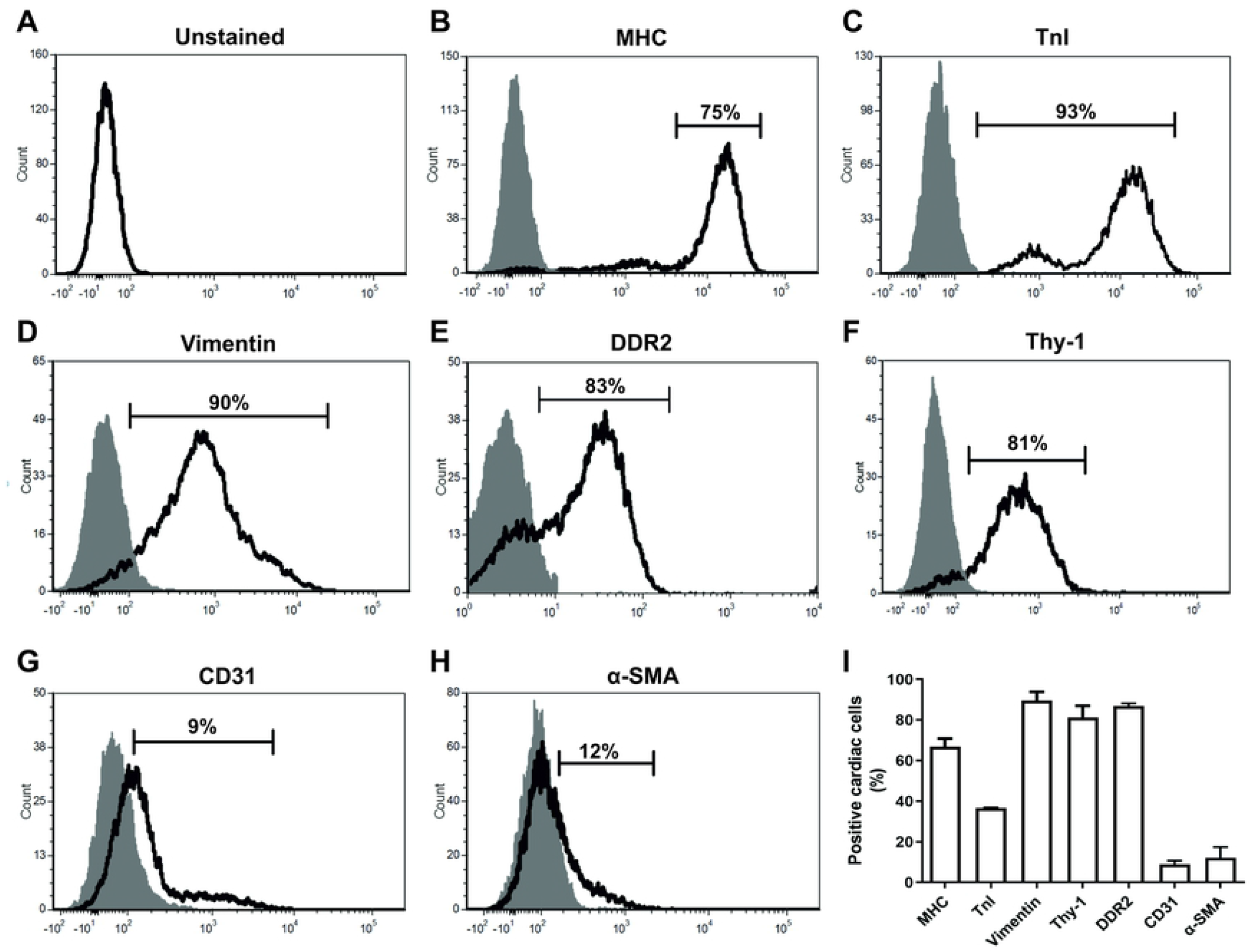
Representative flow cytometry histograms of fetal human heart cells stained for cardiac cell markers. (A) unstained heart cells, (B) MHC, (C) TnI, (D) Vimentin, (E) DDR2, (F) Thy-1, (G) CD31, (H) α-SMA. Grey peaks in flow cytometry histograms represents isotype/negative controls. (I) Histogram showing the percentage of positive cardiac cells for each marker. Error bars represent standard deviation.

Flow cytometry dot plots for TnI and vimentin both revealed two distinct populations of positive cells based on their relative fluorescence intensity (Fig 3). On average, 11% of cells were Vim^High^ (SEM ±1.01) and 78% were Vim^Low^ (SEM ±0.54) (n=7) (Fig 3A and B). On average, 79% of cells were (TnI^High^) (SEM ±2.41) and 19% were TnI^Low^ (SEM ±2.1) (n=14) (Fig 3C and D). High and low expressing populations were also detected for troponin subunit T (TnT) (Supplementary Fig 1). Flow cytometry analysis of cells of the fetal aorta/pulmonary artery, which are absent of cardiomyocytes, expressed TnI at the same fluorescence intensity as the TnI^Low^ cardiac population, confirming this population contains non-cardiomyocyte cells types (Fig 3E).

**Figure 3.**
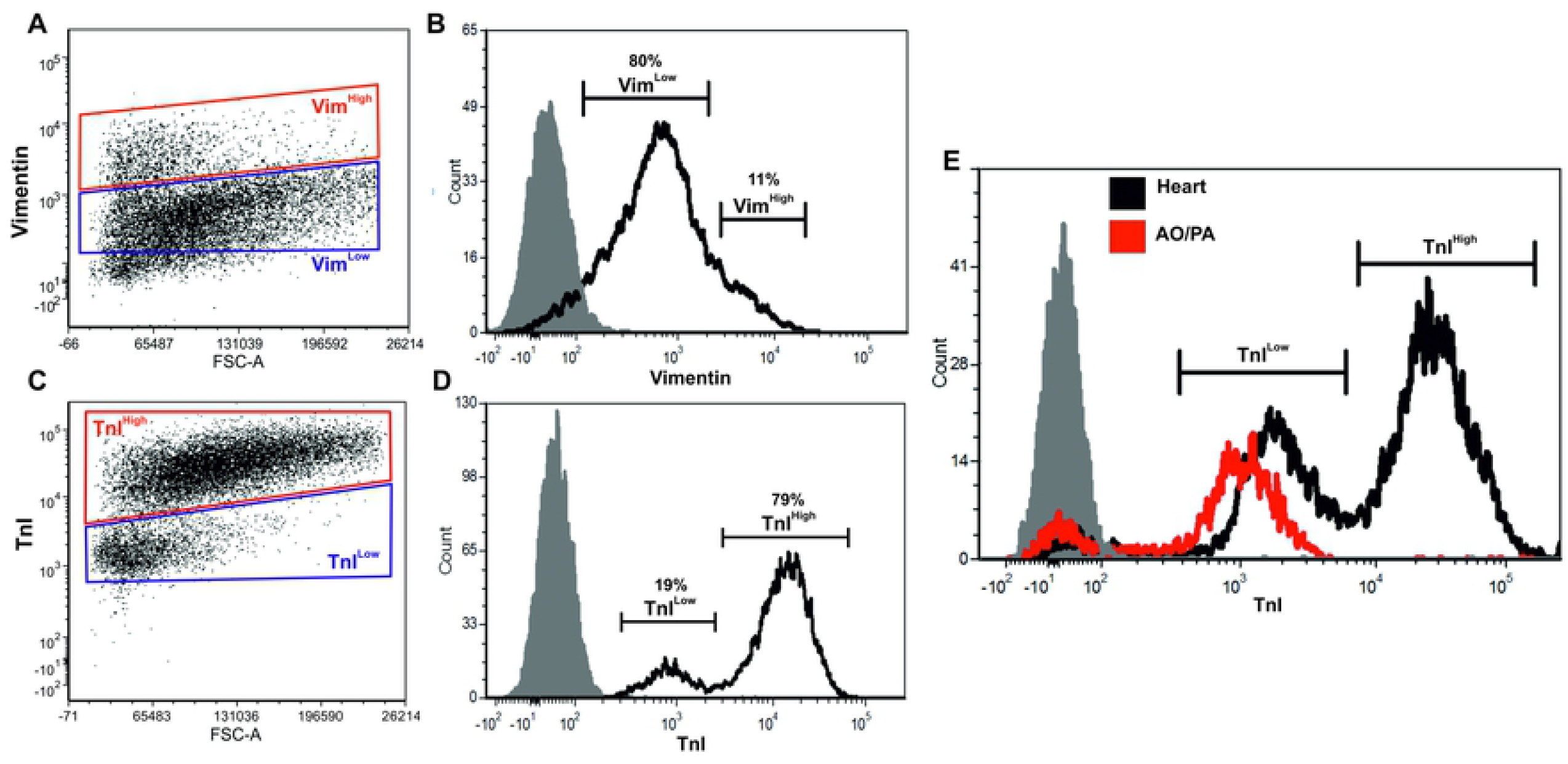
Vimentin and TnI exhibit high and low expressing populations in the human fetal heart. (A) Representative flow cytometry dot plot of vimentin expression in fetal cardiac cells. (B) Representative flow cytometry histogram of vimentin expression in fetal cardiac cells. (C) Representative flow cytometry dot plot of TnI expression in fetal cardiac cells. (B) Representative flow cytometry histogram of TnI expression in fetal cardiac cells. (E) Representative flow cytometry histogram of TnI expression in heart and aorta/pulmonary artery (AO/PA) tissue. Grey peaks in histograms represent isotype controls.

**Supplementary Figure 1.**
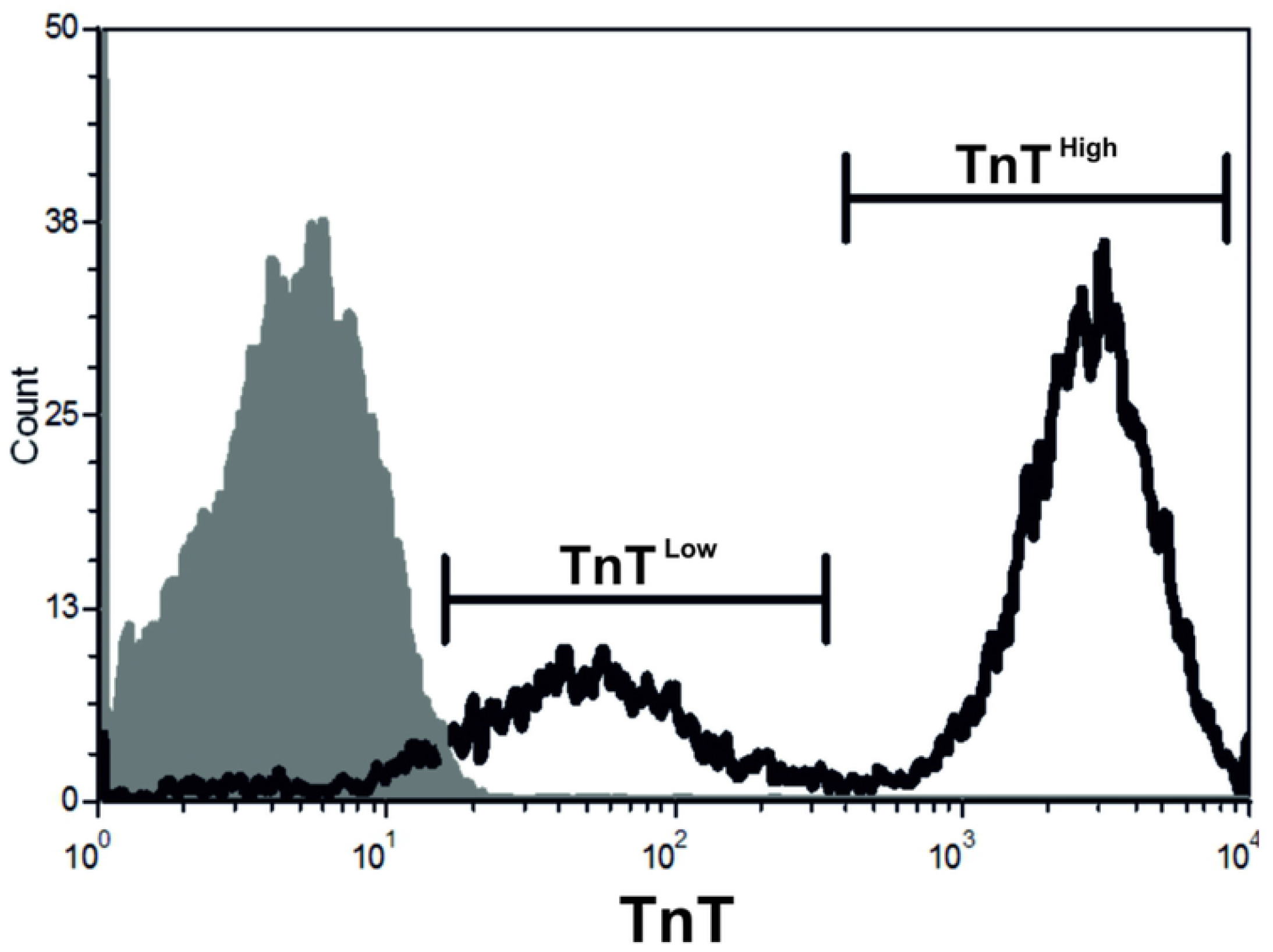
TnT expression in fetal human heart cells. Representative flow cytometry histogram of fetal human heart cells stained for TnT showing two distinct populations of high and low TnT expression. Isotype control shown in grey

### Dual marker flow cytometry and immunohistochemistry of fetal human heart

To interrogate the cellular composition of the two TnI^+^ populations, cardiac cells were co-stained with TnI and MHC and analysed by dual flow cytometry. TnI^High^ cells were also MHC^+^ (80% of total cells, SEM ±2.6) whilst TnI^Low^ cells were MHC^-^ (18% of total cells, SEM ±2.64) (n=5) (Fig 4A). To investigate the identity of Vimentin^+^ populations, cardiac cells were co-stained with vimentin and α-SMA, CD31 or MHC. Vim^High^ and Vim^Low^ cells were then gated (Fig 4B) and plotted against these markers (Fig 4C-F). The majority of Vim^Low^ cells (blue) expressed TnI at higher levels, whilst Vim^High^ cells (red) expressed TnI at lower levels (Fig 4C). On average, 67% of cardiac cells were Vim^Low^/TnI^High^ (SEM ±3.38) and 20% were Vim^High^/TnI^Low^ (SEM ±2.53) (n=4). 18% of cells were α-SMA^+^/Vim^+^ (n=2, SEM ±6.37), with the majority of this population expressing vimentin at lower levels (Fig 4D). 9% of cells were CD31^+^/Vim^+^ (n=8, SEM ±0.84), with the majority of these cells expressing high levels of vimentin (Fig 4E). 79% of cells were MHC^+^/Vim^+^ (n=7, SEM ±1.88), with the majority of this population expressing low levels of vimentin (Fig 4F). Together, these data suggest that fetal cardiomyocytes and cardiac smooth muscle cells express vimentin at lower levels, whilst fetal cardiac endothelial cells express vimentin at higher levels.

**Figure 4.**
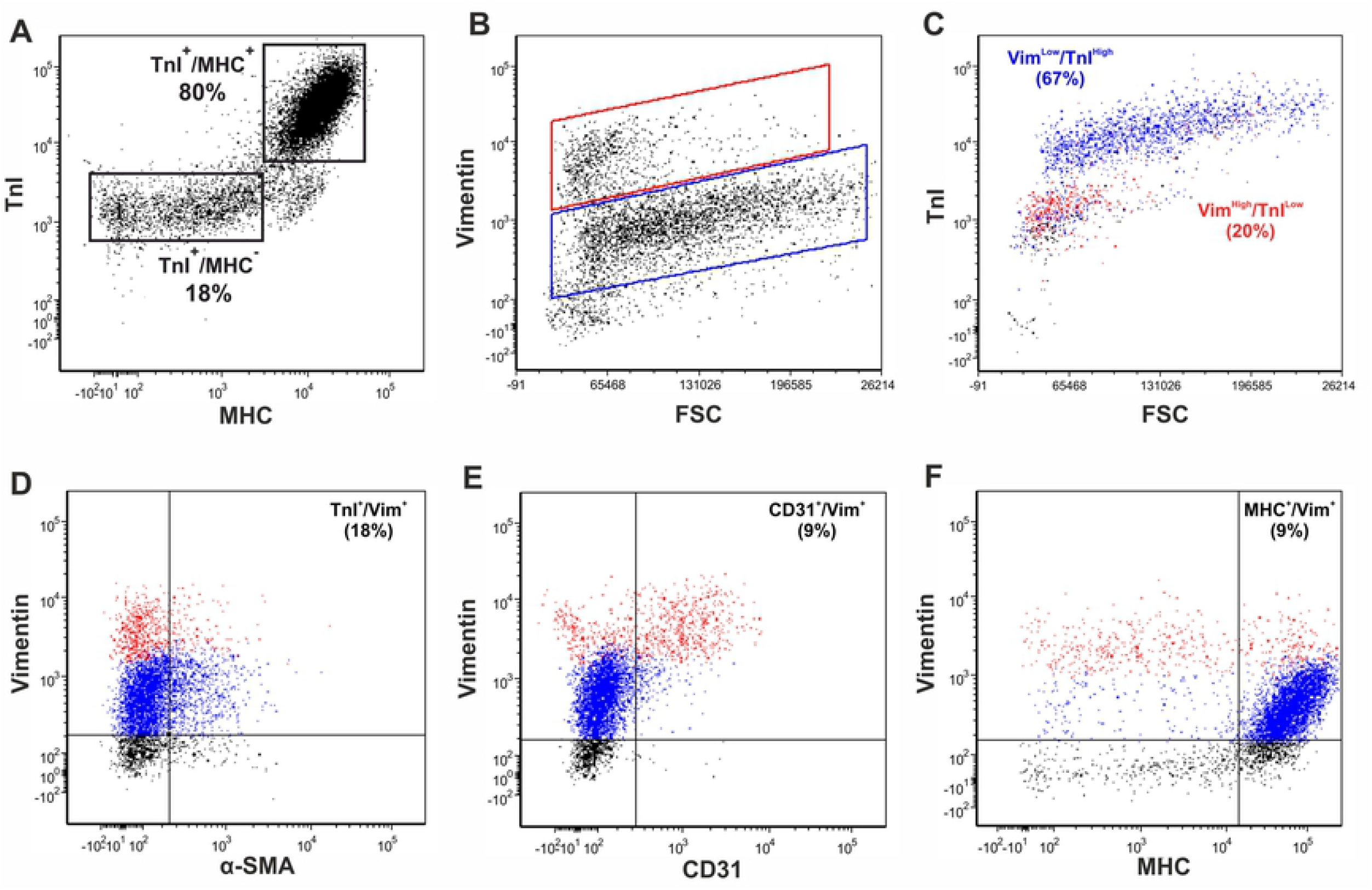
Dual marker flow cytometry of vimentin in human fetal cardiac cell populations. Dual flow cytometry dot plot of TnI and MHC. (B) Gating and colour coding of Vim^High^ (red) and Vim^Low^ (blue) positive cell populations. (C) TnI fluorescence intensity in Vim^High^ and Vim^Low^ cardiac populations. (D) Dual flow cytometry dot plots of vimentin with α-SMA. (E) Dual flow cytometry dot plots of vimentin with CD31. (F) Dual flow cytometry dot plots of vimentin with MHC, populations. Flow cytometry gating based on isotype controls. FSC= forward scatter.

To confirm the flow cytometry data, dual immunohistochemistry was carried out on fetal human heart tissue. Co-expression of TnI and vimentin was seen in the myocardium and the cells of large blood vessels. However, the fluorescence intensity of TnI in the cells lining the blood vessel was much weaker relative to the myocardium, supporting the flow cytometry data that showed high TnI expression in myocytes and low TnI expression in non-myocytes (Fig 5A). Colocalisation of α-SMA and vimentin was observed in coronary vessels (Fig 5B). Similarly, vimentin and CD31 co-stained the endothelial cells lining the cardiac blood vessels (Fig 5C). Co-localisation of vimentin with MHC in the myocardium supported the flow cytometry data that human fetal cardiomyocytes express vimentin (Fig 5D).

**Figure 5.**
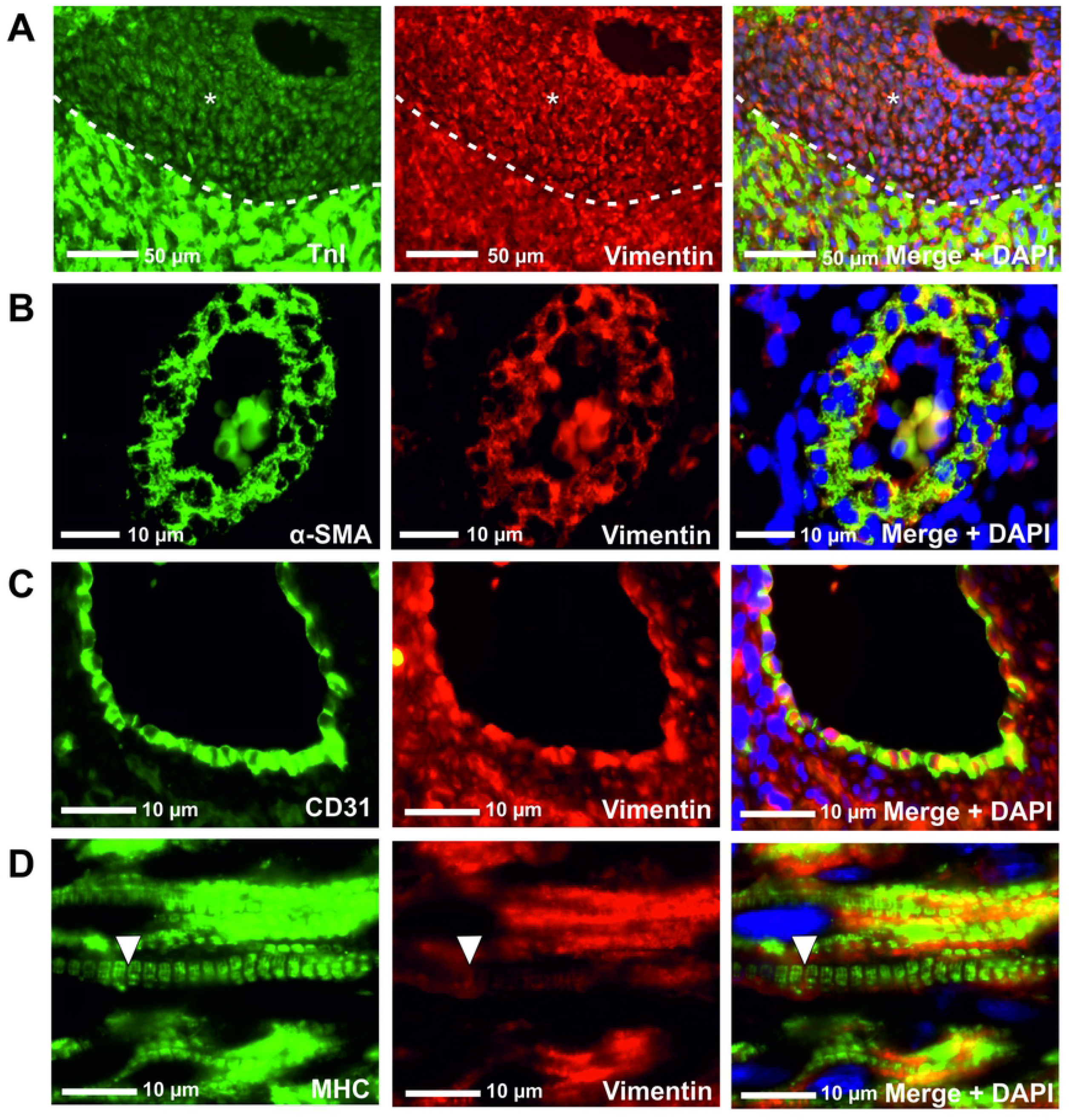
Dual marker immunohistochemistry of vimentin in human fetal cardiac cell populations. Dual immunohistochemistry of vimentin with (A) TnI, (B) α-SMA, (C) CD31 and (D) MHC. Asterisks denote blood vessel wall. Arrows identify sarcomeric structures.

Expression of DDR2 co-localised with α-MHC^+^ cardiomyocytes and α-MHC^-^ cells of the aorta/pulmonary artery vessels (Supplementary Fig 2A). CD31^+^ endothelial cells and α-SMA^+^ smooth muscle cells lining blood vessel also co-stained for DDR2 (Supplementary Fig 2B and C).

**Supplementary Figure 2.**
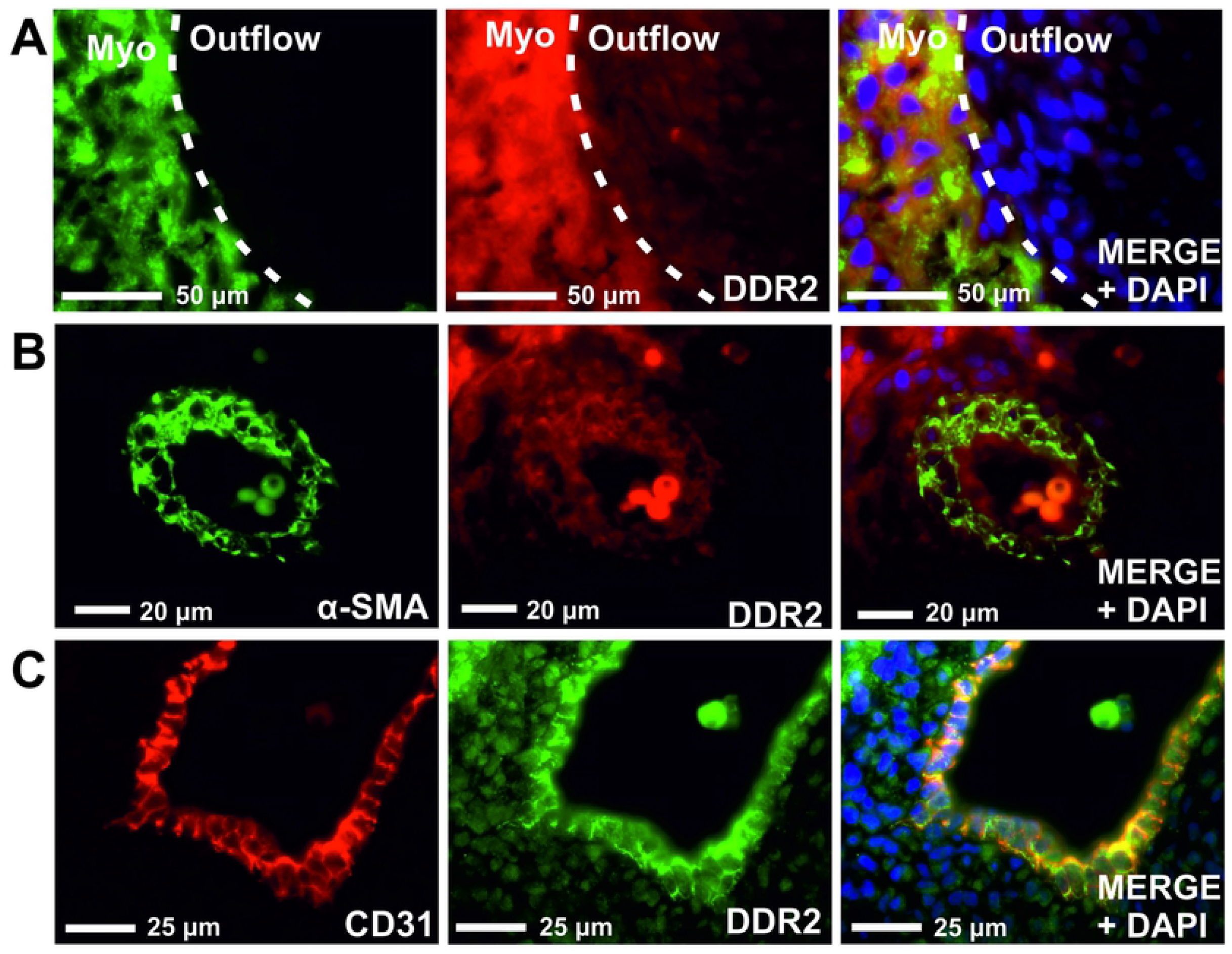
Dual marker immunohistochemistry of DDR2 in human fetal cardiac cell populations. Dual immunohistochemistry of DDR2 with (A) α-MHC, (B) α-SMA and (C) CD31. Myo= myocardium. AO/PA= aorta/pulmonary artery.

### RT-PCR analysis of cardiac cell marker gene expression

To investigate the expression of cardiac cell markers in myocyte and non-myocyte populations, RT-PCR was performed to detect marker transcripts in fetal heart and primary fetal fibroblasts isolated from fetal human ventricular heart tissue, as previously described^18^. Fetal heart tissue expressed *VIM* (*Vimentin), MYH6* (α-*MHC), TNNI3 (cTnI), TNNT2* (*cTnT), PECAM-1 (CD31)* and *α-SMA* (Fig 6A). Cardiac fibroblasts were enriched for VIM and α-SMA (Fig 6B). *TNNI3 and TNNT2* transcripts were also detected in cardiac fibroblasts, confirming the expression of these markers in non-myocyte populations. α-*MHC* mRNA expression was not detected in cardiac fibroblasts, confirming that expression of *TNNI3* and *TNNT2* was not due to cardiomyocyte contamination. The absence of CD31 mRNA expression also confirmed that the isolated fibroblasts were absent of endothelial cells.

**Figure 6.**
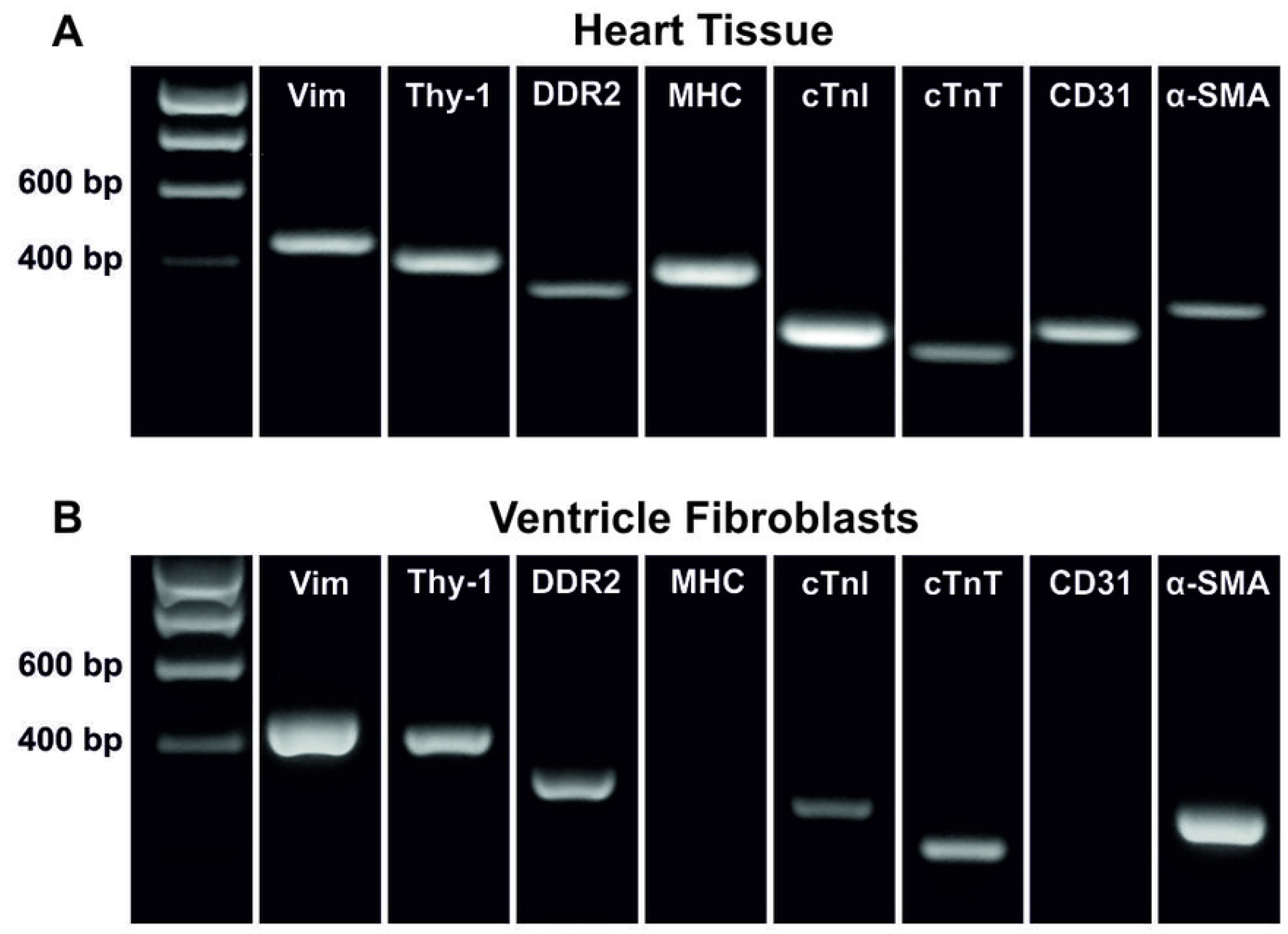
Gene expression analysis of cardiac markers in human fetal heart and ventricular fibroblasts. (A) RT-PCR gene expression analysis of previously described cell markers in fetal human heart. (B) RT-PCR gene expression analysis of cell markers in isolated fetal human cardiac fibroblasts. Non-contiguous gel lanes are demarcated by vertical white lines.

### Marker protein expression in fetal ventricular fibroblasts

Cultured ventricular fibroblasts were stained for Vimentin, TnI and α-SMA and analysed by flow cytometry and immunocytochemistry. All three proteins were enriched in ventricular fibroblasts (>96% positive) (Fig 7). This data further supports the expression of TnI in non-myocyte cells of the fetal heart. Interestingly, the staining pattern of TnI in these cells is distinct from that observed in mature cardiomyocytes, with a lack of sarcomeric organisation.

**Figure 7.**
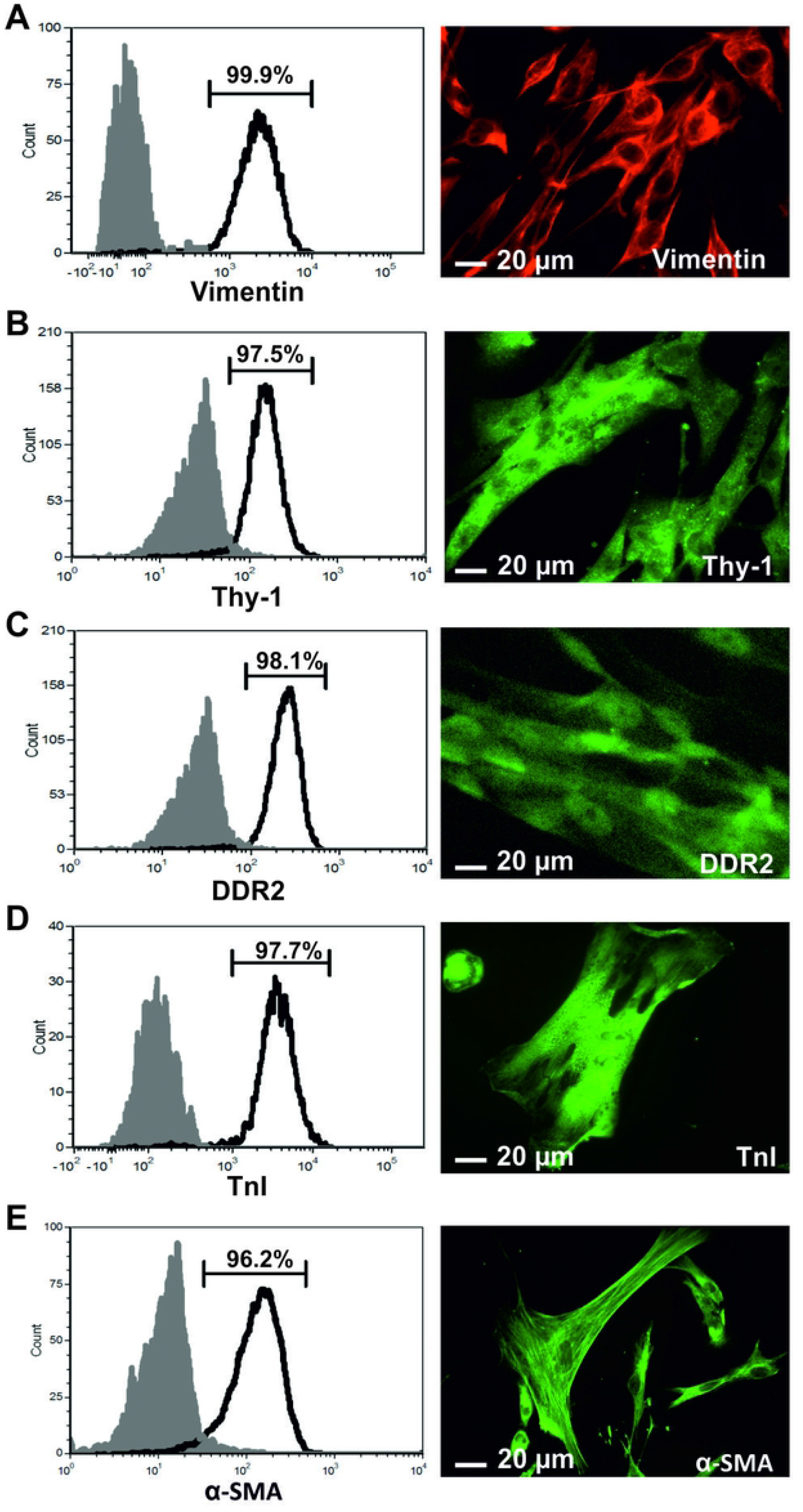
Marker protein expression in fetal ventricular fibroblasts. Representative flow cytometry and immunocytochemistry analyses of (A) vimentin (B) Thy-1 (C), DDR2 (D), TnI and (E) α-SMA Primary cultures fetal ventricular fibroblasts (n=3). Isotype controls in grey.

## Discussion

This study aimed to investigate the expression and specificity of commonly used cardiac cell markers in the early developing human heart. Immunohistochemical analyses of the cardiomyocyte markers α-MHC and cTnI showed defined sarcomeric staining in the myocardium of fetal heart tissue confirming their expression in cardiomyocytes. However, flow cytometry analyses showed a greater percentage of cardiac cells expressing cTnI (93%) compared to α-MHC (79%), suggesting cells other than cardiomyocytes may be expressing cTnI. Flow cytometry, immunocytochemistry and RT-PCR analyses were able to confirm the expression of cTnI at lower levels in non-myocytes and higher levels in cardiomyocytes. This is supported by recent findings from Cui *et al*.,^28^ identifying positive immunostaining of troponin T in cardiac fibroblasts from 17 gestational week fetal cardiac tissue. Furthermore, previous studies have shown the expression of other contractile proteins, including actin, myosin and tropomyosin, in non-muscle cells, which suggests that cardiac troponin may also have non-muscle-specific isoforms ^29-31^. In this study, the non-striated pattern of cTnI expression in cardiac fibroblasts confirmed a lack of sarcomeric organisation, suggesting a role of troponins in non-muscle cells distinct from that of cardiomyocytes and skeletal muscle cells. As such, this data suggests troponin proteins are not specific markers of cardiomyocytes in the developing human heart.

The mesenchymal marker vimentin is a commonly used fibroblast marker; however, we showed approximately 90% of fetal cardiac cells are positive for vimentin. We confirmed that endothelial cells express high levels of vimentin, as previously described^15^. Whilst our data confirmed the enrichment of vimentin in cultured cardiac fibroblasts, flow cytometry analysis of fetal heart cells showed expression of vimentin at low levels in α-MHC^+^ cardiomyocytes, with 79% of cardiac cells having α-MHC^+^/Vim^Low^ marker profile. This is consistent with a previous study that identified weak expression of vimentin in some cardiomyocytes of 9-14 pcw human hearts and demonstrated that with increasing fetal age vimentin expression in cardiomyocytes decreases and desmin expression increases ^32^. The lack of data identifying vimentin expression in adult cardiomyocytes further suggests that vimentin expression may be a property of cardioymyocytes of the developing heart only. The collagen receptor DDR2 has been identified as a more specific marker of cardiac fibroblasts^33^, however, our results confirmed its expression in cardiomyocytes, endothelial cells and smooth muscle cells of the developing heart.

Together, our results offer an estimate of the relative cellular composition of the human fetal heart. We consider the marker profile ‘α-MHC^+^/cTnI^High^/Vim^Low^’ to represent the fetal human cardiomyocyte population, which we estimate to comprise 75-80% of total cardiac cells. The non-cardiomyocyte population (primarily composed of fibroblasts, smooth muscle cells and endothelial cells) comprises 20-25% of total cardiac cells and exhibit an α-MHC^-^/cTnI^Low^/Vim^High^ marker profile, of which approximately 10% are endothelial cells (CD31^+^/Vim^High^). Due to the lack of a fibroblast-specific marker, it is difficult to accurately determine the percentage of fibroblasts and smooth muscle cells, although it is likely in the region of 5-15%. Of note, we found 11% of cells were Vim^+^/α-SMA^+^ which may represent the combined fibroblast and smooth muscle cell population or in fact a myofibroblast population. The common lineage of smooth muscle cells and fibroblasts predicts that during early fetal development, discrimination between these cell types may be difficult if the phenotypes have yet to become distinct ^28,34^.

These results indicate a fetal cellular composition distinct from that of the adult heart, which has been estimated to be comprised of 30-40% cardiomyocytes, despite the cells occupying three quarters of normal myocardial volume. Our results suggest that in the early developing heart, cardiomyocytes are the predominant cell type, comprising around 75% of total cells. This figure is likely reflective of the proliferative nature of fetal cardiomyocytes relative to adult cardiomyocytes. Mitosis of differentiated cardiomyocytes in the developing heart is well documented and is responsible for cardiac morphogenesis and organogenesis in utero^35^, however, from two weeks after birth, cardiomyocyte proliferation is significantly reduced in the mammalian heart as these cells enter cell cycle arrest, with continued cardiac growth reliant of hypertrophy of pre-existing cardiomyocytes ^36,37^.

The results from our study suggest the marker profiles of fetal cardiac cells are distinct from that of adult cells. The phenotypic similarities between the mid-gestation human fetal heart and pluripotent stem cell-derived cardiomyocytes suggests our data could be useful when purifying and characterising these cells. Furthermore, the marker profiles identified could potentially be used for further studies to determine how the ratio of cardiomyocytes to non-crdiomyocytes changes throughout fetal development.

## Acknowledgments

This work was funded by the MRC DTA Capacity Building Studentship G1000406. The human embryonic and fetal material was provided by the Joint MRC/Wellcome Trust grant #099175/Z/12/Z Human Developmental Biology Resource (HDBR)

## Notes

### Competing Interest Statement

The authors have declared no competing interest.

## References

1 Vliegen, H. W., Laarsem A. V.D, Cornelisse, C.J. Eulderink, F. Myocardial changes in pressure overload-induced left ventricular hypertrophy. A study on tissue composition, polyploidization an multinucleation. European Heart Journal 12, 488–494 (1991).

2 Camelliti, P., Borg, T. K. & Kohl, P. Structural and functional characterisation of cardiac fibroblasts. Cardiovascular research 65, 40–51, doi:10.1016/j.cardiores.2004.08.020 (2005).

3 Moore, G. W., Hutchins, G. M., Bulkley, B. H., Tseng, J. S. & Ki, P. F. Constituents of the human ventricular myocardium: connective tissue hyperplasia accompanying muscular hypertrophy. Am Heart J 100, 610–616 (1980).

4 Pinto, A. R. et al. Revisiting Cardiac Cellular Composition. Circulation research 118, 400–409, doi:10.1161/CIRCRESAHA.115.307778 (2016).

5 Zhou, P. & Pu, W. T. Recounting Cardiac Cellular Composition. Circ Res 118, 368–370, doi:10.1161/CIRCRESAHA.116.308139 (2016).

6 Denning, C. et al. Cardiomyocytes from human pluripotent stem cells: From laboratory curiosity to industrial biomedical platform. Biochimica et biophysica acta, doi:10.1016/j.bbamcr.2015.10.014 (2015).

7 Souders, C. A., Bowers, S. L. & Baudino, T. A. Cardiac fibroblast: the renaissance cell. Circulation research 105, 1164–1176, doi:10.1161/CIRCRESAHA.109.209809 (2009).

8 Ponten, A. et al. FACS-based isolation, propagation and characterization of mouse embryonic cardiomyocytes based on VCAM-1 surface marker expression. PloS one 8, e82403, doi:10.1371/journal.pone.0082403 (2013).

9 Banerjee, I., Fuseler, J. W., Price, R. L., Borg, T. K. & Baudino, T. A. Determination of cell types and numbers during cardiac development in the neonatal and adult rat and mouse. Am J Cardiovasc Dis 293, H1883–H1891, doi:10.1152/ajpheart.00514.2007.-Cardiac (2007).

10 Goldsmith, E. C., Zhang, X., Watson, J., Hastings, J. & Potts, J. D. The collagen receptor DDR2 is expressed during early cardiac development. Anat Rec (Hoboken) 293, 762–769, doi:10.1002/ar.20922 (2010).

11 Goldsmith, E. C. et al. Organization of fibroblasts in the heart. Developmental Dynamics 230, 787–794, doi:10.1002/dvdy.20095 (2004).

12 Shyu, K. G., Chao, Y. M., Wang, B. W. & Kuan, P. Regulation of discoidin domain receptor 2 by cyclic mechanical stretch in cultured rat vascular smooth muscle cells. Hypertension 46, 614–621, doi:10.1161/01.HYP.0000175811.79863.e2 (2005).

13 Morales, M. O., Price, R. L. & Goldsmith, E. C. Expression of Discoidin Domain Receptor 2 (DDR2) in the developing heart. Microscopy and microanalysis: the official journal of Microscopy Society of America, Microbeam Analysis Society, Microscopical Society of Canada 11, 260–267, doi:10.1017/S1431927605050518 (2005).

14 Wang, R., Li, Q. & Tang, D. D. Role of vimentin in smooth muscle force development. Am J Physiol Cell Physiol 291, C483–C489 (2011).

15 Dave, J. M. & Bayless, K. J. Vimentin as an integral regulator of cell adhesion and endothelial sprouting. Microcirculation 21, 333–344, doi:10.1111/micc.12111 (2014).

16 Hudon-David, F., Bouzeghrane, F., Couture, P. & Thibault, G. Thy-1 expression by cardiac fibroblasts: lack of association with myofibroblast contractile markers. Journal of molecular and cellular cardiology 42, 991–1000, doi:10.1016/j.yjmcc.2007.02.009 (2007).

17 Baum, J. & Duffy, H. S. Fibroblasts and myofibroblasts: what are we talking about? Journal of cardiovascular pharmacology 57, 376–379, doi:10.1097/FJC.0b013e3182116e39 (2011).

18 Ieda, M. et al. Direct reprogramming of fibroblasts into functional cardiomyocytes by defined factors. Cell 142, 375–386, doi:10.1016/j.cell.2010.07.002 (2010).

19 Rege, T. A. & Hagood, J. S. Thy-1 as a regulator of cell-cell and cell-matrix interactions in axon regeneration, apoptosis, adhesion, migration, cancer, and fibrosis. FASEB journal: official publication of the Federation of American Societies for Experimental Biology 20, 1045–1054, doi:10.1096/fj.05-5460rev (2006).

20 Craig, W., Kay, R., Cutler, R. L. & Lansdorp, P. M. Expression of Thy-1 on human hematopoietic progenitor cells. J. Exp. Med. 177, 1331–1342 (1993).

21 Wetzel, A. et al. Human Thy-1 (CD90) on Activated Endothelial Cells Is a Counterreceptor for the Leukocyte Integrin Mac-1 (CD11b/CD18). The Journal of Immunology 172, 3850–3859, doi:10.4049/jimmunol.172.6.3850 (2004).

22 Kong, P., Christia, P., Saxena, A., Su, Y. & Frangogiannis, N. G. Lack of specificity of fibroblast-specific protein 1 in cardiac remodeling and fibrosis. American journal of physiology. Heart and circulatory physiology 305, H1363–1372, doi:10.1152/ajpheart.00395.2013 (2013).

23 Protze, S. et al. A new approach to transcription factor screening for reprogramming of fibroblasts to cardiomyocyte-like cells. Journal of molecular and cellular cardiology 53, 323–332, doi:10.1016/j.yjmcc.2012.04.010 (2012).

24 Qian, L., Berry, E. C., Fu, J. D., Ieda, M. & Srivastava, D. Reprogramming of mouse fibroblasts into cardiomyocyte-like cells in vitro. Nature protocols 8, 1204–1215, doi:10.1038/nprot.2013.067 (2013).

25 Molkentin, J. D., Jobe, S. M. & Markham, B. E. α-myosin heavy chain gene regulation: delineation and characterization of the cardiac muscle-specific enhancer and muscle-specific promoter. Journal of molecular and cellular cardiology 28, 1211–1225 (1996).

26 Saunders, V., Dewing, J. M., Sanchez-Elsner, T. & Wilson, D. I. Expression and localisation of thymosin beta-4 in the developing human early fetal heart. PLOS ONE 13, e0207248, doi:10.1371/journal.pone.0207248 (2018).

27 Baudino, T. A., Carver, W., Giles, W. & Borg, T. K. Cardiac fibroblasts: friend or foe? American journal of physiology. Heart and circulatory physiology 291, H1015–1026, doi:10.1152/ajpheart.00023.2006 (2006).

28 Cui, Y. et al. Single-Cell Transcriptome Analysis Maps the Developmental Track of the Human Heart. Cell Rep 26, 1934–1950 e1935, doi:10.1016/j.celrep.2019.01.079 (2019).

29 Ju, Y. et al. Troponin T3 expression in skeletal and smooth muscle is required for growth and postnatal survival: characterization of Tnnt3(tm2a(KOMP)Wtsi) mice. Genesis 51, 667–675, doi:10.1002/dvg.22407 (2013).

30 Kajioka, S. et al. Endogenous cardiac troponin T modulates Ca(2+)-mediated smooth muscle contraction. Scientific reports 2, 979, doi:10.1038/srep00979 (2012).

31 Gahlmann, R., Wade, R., Gunning, P. & Kedes, L. Differential expression of slow and fast skeletal muscle troponin C: Slow skeletal muscle troponin C is expressed in human fibroblasts. Journal of Molecular Biology 201, 379–391 (1988).

32 Kim, H. D. Expression of intermediate filament Desmin and Vimentin in the human fetal heart. Anat Rec 246, 271–278 (1996).

33 Tarbit, E., Singh, I., Peart, J. N. & Rose’Meyer, R.B. Biomarkers for the identification of cardiac fibroblast and myofibroblast cells. Heart Fail Rev 24, 1–15, doi:10.1007/s10741-018-9720-1 (2019).

34 Rinkevich, Y. et al. Identification and prospective isolation of a mesothelial precursor lineage giving rise to smooth muscle cells and fibroblasts for mammalian internal organs, and their vasculature. Nature Cell Biology 14, 1251–1260 (2012).

35 Yutzey, K. E. Cardiomyocyte Proliferation: Teaching an Old Dogma New Tricks. Circ Res 120, 627–629, doi:10.1161/CIRCRESAHA.116.310058 (2017).

36 Leri, A., Rota, M., Pasqualini, F. S., Goichberg, P. & Anversa, P. Origin of cardiomyocytes in the adult heart. Circulation research 116, 150–166, doi:10.1161/CIRCRESAHA.116.303595 (2015).

37 Kikuchi, K. & Poss, K. D. Cardiac regenerative capacity and mechanisms. Annual review of cell and developmental biology 28, 719–741, doi:10.1146/annurev-cellbio-101011-155739 (2012).

